# Recurrent processing drives experience-dependent plasticity for perceptual decisions

**DOI:** 10.1101/2020.04.07.030510

**Authors:** Ke Jia, Elisa Zamboni, Valentin Kemper, Catarina Rua, Nuno Reis Goncalves, Adrian Ka Tsun Ng, Christopher T. Rodgers, Guy Williams, Rainer Goebel, Zoe Kourtzi

## Abstract

Learning and experience are critical for translating ambiguous sensory information from our environments to perceptual decisions. Yet, evidence on how training molds the adult human brain remains controversial, as fMRI at standard resolution does not allow us to discern the finer-scale mechanisms that underlie sensory plasticity. Here, we combine ultra-high field (7T) functional imaging at sub-millimetre resolution with orientation discrimination training to interrogate experience-dependent plasticity across cortical depths. Our results provide evidence for recurrent plasticity, by contrast to sensory encoding vs. feedback mechanisms. We demonstrate that learning alters orientation-specific representations in superficial rather than middle V1 layers, suggesting changes in read-out rather than input signals. Further, learning increases feedforward rather than feedback layer-to-layer connectivity in occipito-parietal regions, suggesting that sensory plasticity gates perceptual decisions. Our findings propose finer-scale plasticity mechanisms that re-weight sensory signals to inform improved decisions, bridging the gap between micro- and macro-circuits of experience-dependent plasticity.

## Introduction

Understanding the world around us depends on the brain resolving ambiguous information from our senses to inform our decisions and actions. The brain learns to interpret sensory signals by using past experience to optimize perceptual judgments. Although this experience-dependent brain plasticity is most evident during development, training in adulthood is shown to improve visual recognition skills and alter the brain’s function and connectivity^1–4^. Yet, evidence on how practice molds the adult human brain and results in improved decisions remains controversial.

The neural computations that mediate experience-dependent plasticity are highly debated (e.g. ^1,5,6^). Results from behavioral training studies focusing on visual discrimination tasks (i.e. perceptual learning tasks) have long been understood to suggest that plasticity occurs at early stages of sensory processing (i.e. primary visual cortex), as learning was shown to be specific to the trained stimulus features (e.g. ^7,8^). Neurophysiological recordings^9,10^ and human imaging data ^11–13^ support this early neural locus hypothesis by showing that training changes neural responses in primary visual cortex, implying that learning alters stimulus encoding. By contrast, other studies have shown that learning alters processing in higher visual areas^14,15^ and regions involved in decision-making^6,16^. This suggests that learning changes the read-out of sensory information^5^ from higher cortical areas rather than sensory encoding in visual cortex, with changes in activity in early visual cortex reflecting feedback processes^17^.

Here, we capitalize on recent advances in brain imaging technology (i.e. Ultra-High Field: UHF imaging) that allow us to interrogate brain computations at a finer scale than that afforded by standard fMRI techniques^18^. UHF imaging affords the sub-millimetre resolution necessary to examine fMRI signals across cortical laminar layers that are known to be associated with dissociable brain computations (Figure 1). In particular, sensory input is known to enter the cortex from the thalamus at the level of the middle layer (layer 4) and output information is fed forward from the superficial layers (layer 2/3), while feedback information is exchanged primarily between deeper (layer 5/6) and superficial layers^19–22^. Further, horizontal connections across V1 columns are known to predominantly terminate in superficial layers^23,24^ and suggested to support recurrent processing within visual cortex^25,26^. Neurophysiological studies have shown that this micro-circuit is involved in a range of visual recognition^26^ and attention^27^ tasks. Recent laminar fMRI studies provide evidence for the involvement of this circuit in the context of sensory processing^28^ and visual attention^29^.

**Figure 1.**
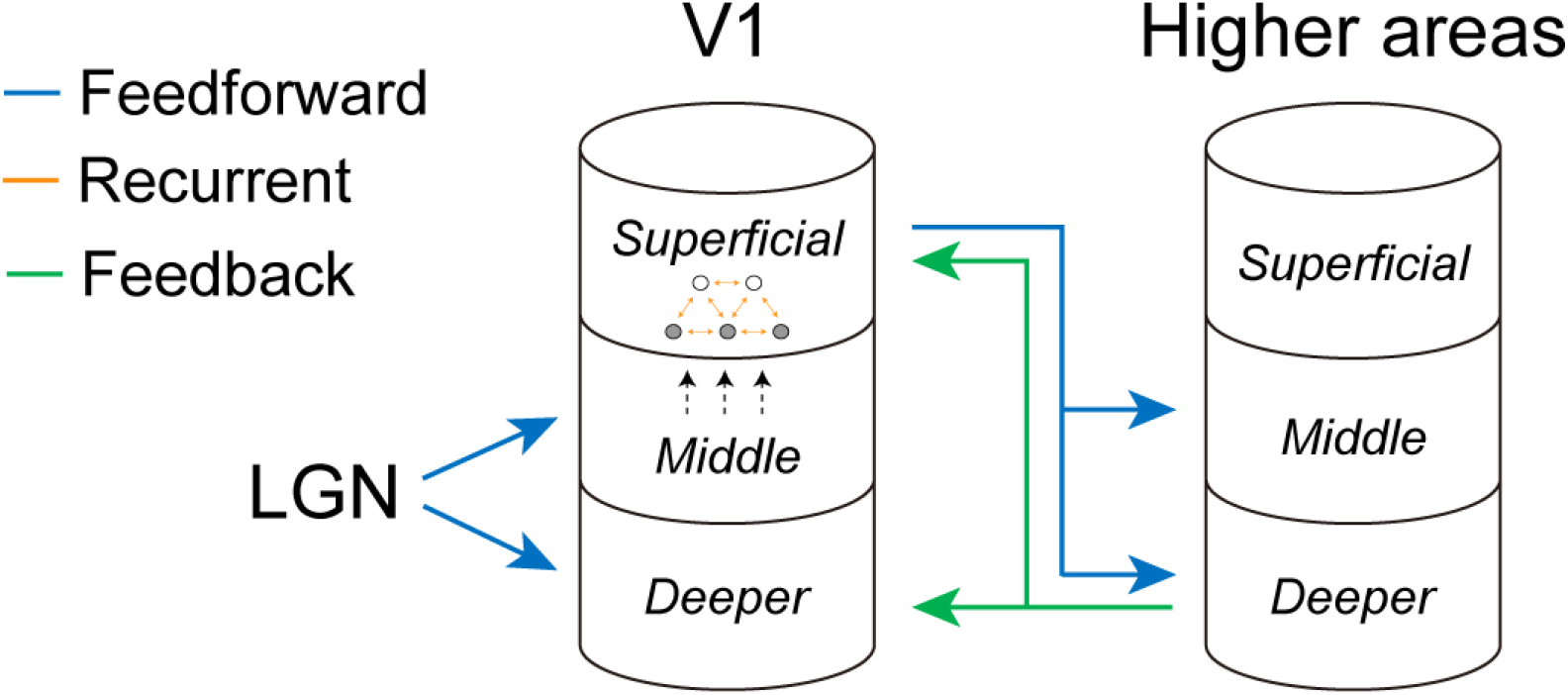
Laminar brain circuits. Schematic representation of the hypotheses tested, training may modify: a) feedforward processing (blue arrows) between LGN and V1; b) recurrent processing via horizontal connections within the visual cortex (horizontal connections (indicated by orange arrows) between excitatory (open circles) and inhibitory (filled circles) neurons adapted from Teich et al. (2003) and Schwabe et al. (2005)); c) feedback processing (green arrows) between V1 and higher areas (i.e., V2, V3, V4, IPS) based on known anatomical circuits.

We combine 7T laminar fMRI (i.e. before and after training) with training on an orientation discrimination task, to test whether learning modifies: a) encoding of sensory input in middle V1 layers, b) recurrent processing in superficial layers that alters the sensory read-out, c) feedback processing in deeper or superficial V1 layers from higher decision-related regions (i.e. intra-parietal cortex, IPS). Using multi-voxel pattern classification analysis (MVPA) across cortical depths, we demonstrate learning-dependent changes in visual representations that are specific to superficial rather than middle or deeper V1 layers, suggesting that learning modifies read-out rather than input signals in visual cortex. Further, we show enhanced feedforward connectivity between superficial layers in visual cortex and middle layers of posterior parietal cortex, rather than feedback connectivity between deeper layers in these areas. Our findings offer an alternate proposal to the ongoing debate focusing on encoding vs. feedback mechanisms of experience-dependent plasticity: learning is implemented by recurrent computations that alter read-out rather than sensory encoding in visual cortex and feedforward processing from sensory to decision related areas.

## Results

### Learning-dependent changes in perceptual discrimination

We trained participants (N = 15; data from two participants were excluded due to excessive head movement and technical issues during acquisition) on an orientation discrimination task^9,12^ for five consecutive days and tested their performance on the same task during fMRI scanning before and after training (Figure 2A&B). Participants’ discrimination performance improved during training (Figure 2C), as indicated by a significant decrease in threshold (~79.4%) performance (paired t-test, mean threshold on day 1 vs. day 5: t(12) = 8.108, *p* < 0.001).

**Figure 2.**
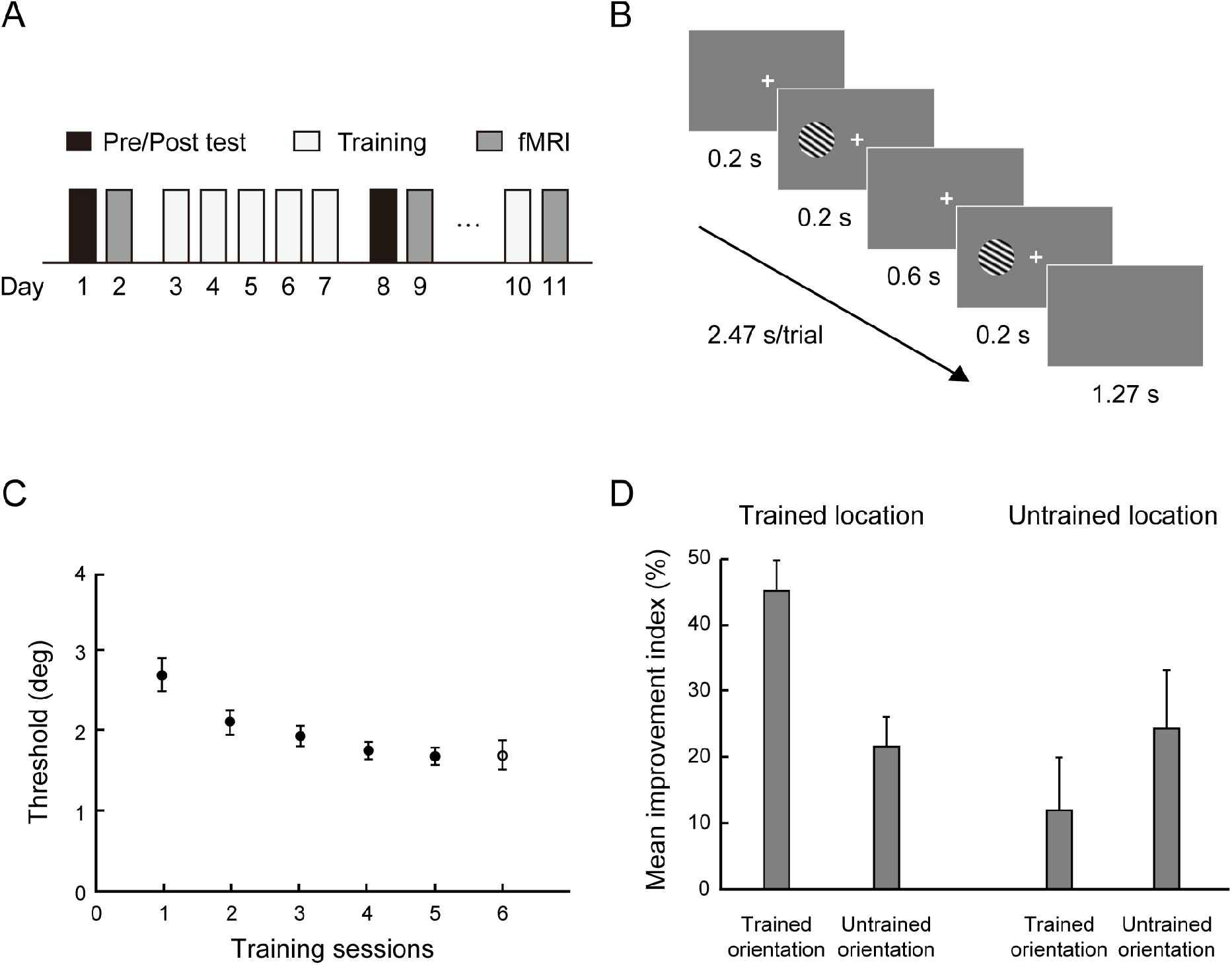
Experimental design, task and behavioral results. (A) Experimental design. Participants were trained on an orientation discrimination task with feedback for five consecutive days. Before and after training, we measured participant’s performance on the same task without feedback in the lab and during scanning. (B) Orientation discrimination task. For each trial, participants were asked to report whether the second grating was tilted clockwise or counterclockwise relative to the first grating. (C) Mean performance across participants at 79.4% threshold for the training (filled circles) and the control (open circle) sessions. (D) Mean improvement index (MPI = (pre-test threshold – post-test threshold) / pre-test threshold × 100%) showed learning specificity for the trained compared to the untrained orientation presented at the trained vs. untrained location. A two-way repeated measures ANOVA on MPI (orientation × location) showed a significant interaction (F(1,12) = 14.847, *p* = 0.002). Post-hoc comparisons showed significantly higher improvement for the trained than the untrained (t(12) = 5.564, *p* < 0.001) orientation at the trained location. In contrast, no significant differences were observed between the trained and the untrained (t(12) = −1.608, *p* = 0.134) orientations at the untrained location. Error bars indicate standard error of the mean across participants.

To determine the specificity of this learning, we measured participants’ discrimination threshold for two different orientations (i.e., trained vs. untrained orientations that corresponded to 55° or 125°) at two different locations (i.e., trained vs. untrained location that corresponded to the left or right visual field) before and after training. Our results showed that behavioral improvement due to training was stronger for the trained orientation and location (Figure 2D, Figure S1), consistent with previous studies^8^ showing specificity to the trained stimulus features. In particular, we observed a significant orientation × location × session interaction (repeated measures ANOVA, F(1,12) = 11.858, *p* = 0.005) and a significant orientation × session interaction (F(1,12) = 21.551, *p* = 0.001) at the trained, but not the untrained (F(1,12) = 3.093, *p* = 0.104) location. As suggested by recent work^7^, it is likely that this learning specificity is due to prolonged training near threshold (i.e. employing a single staircase^30^), in contrast to supra-threshold training that has been suggested to enhance transfer and higher-level learning^31^.

### Learning-dependent changes across cortical depth in the visual cortex

To test whether learning alters orientation representations across cortical depth in the visual cortex, we segmented the visual areas and assigned voxels to three layers (superficial, middle, deeper) using an equi-volume approach (*see Methods, Anatomical data analyses for details*; Figure 3A-D). We used MVPA to discern orientation-specific fMRI signals and test for differences in these signals before vs. after training across layers. In particular, we tested whether linear classifiers that were trained on fMRI signals from multi-voxel patterns in different V1 layers (superficial, middle, deeper) discriminated between: a) trained (55° or 125°) vs. control (vertical) orientations, b) untrained (125° or 55°) vs. control (vertical) orientations. Classifying each of the reference orientations (55° or 125°) from the control (vertical) orientation allowed us to test learning-dependent changes separately for the trained vs. untrained orientations. We hypothesized that higher MVPA accuracy after training for the trained vs. control orientation classification than the untrained vs. control orientation classification would indicate orientation-specific learning-dependent plasticity. Note that, as both the trained and untrained orientation differed equally from the control orientation (~55°), accuracy differences between these classification tasks could not be attributed to stimulus differences.

**Figure 3.**
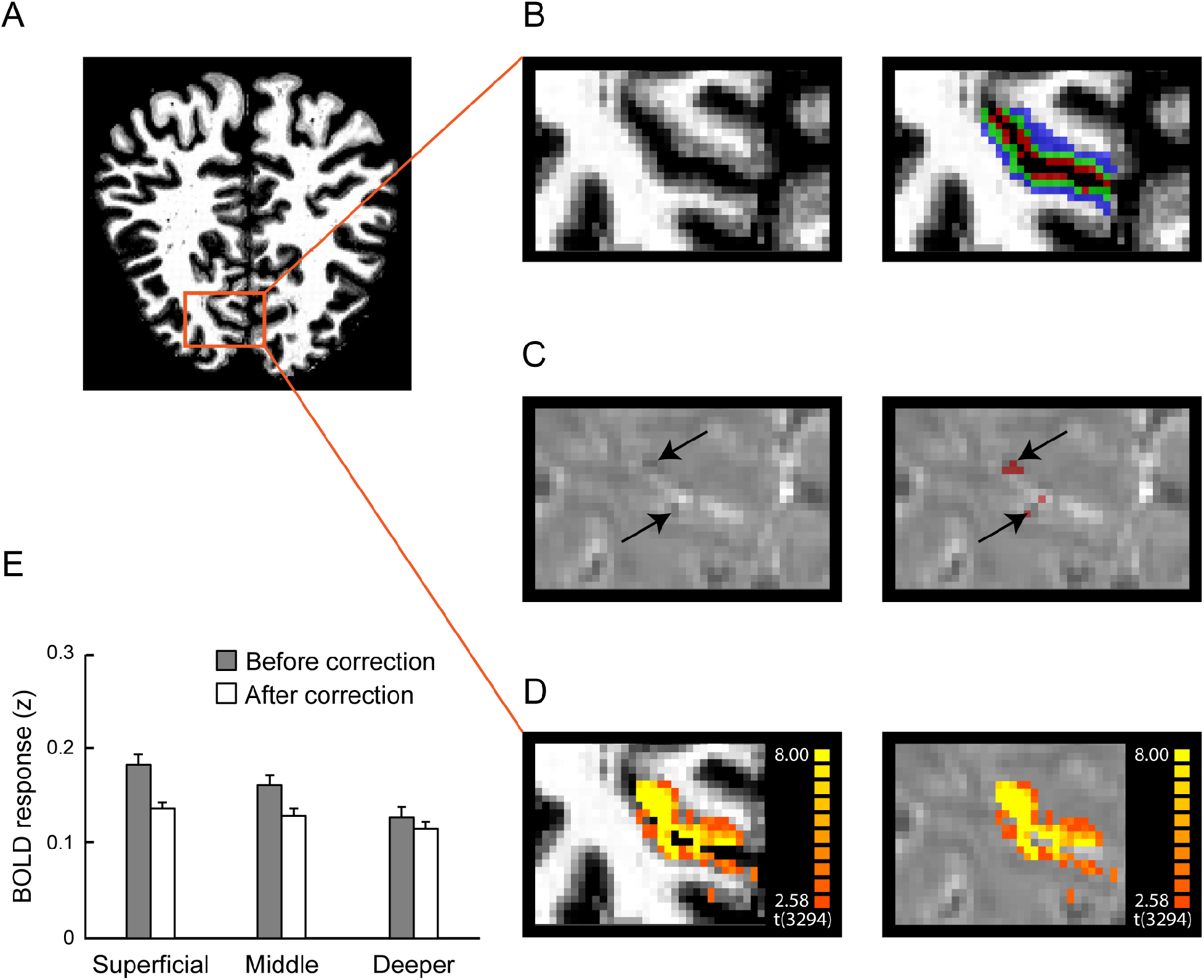
fMRI layer definition and vascular correction. (A) Coronal view of the anatomical image of a sample participant. Red insert indicates region of interest in visual cortex. (B) Layers definition map overlaid on an anatomical image (blue: deeper layers, green: middle layers, and red: superficial layers). (C) Voxels confounded by vasculature-related effects (highlighted by arrows and in red) overlaid on functional images. (D) BOLD activation map (stimulus vs. fixation) overlaid on the anatomical (left panel) and functional data (right panel). (E) Mean normalized BOLD in V1 before and after correction for vasculature-related effects, showing reduced superficial bias after correction. Error bars indicate standard error of the mean across participants. We observed significant interactions (pre-test session: F(2,24) = 50.961, *p* < 0.001, post-test session: F(2,24) = 36.887, *p* < 0.001) between layer (superficial, middle, deeper) and BOLD signal (before vs. after correction). The stronger BOLD decrease in upper (i.e. superficial, middle) than deeper layers after correction suggests that our approach for correcting vasculature-related effects controlled substantially for the superficial bias.

Our results showed learning-dependent changes (i.e. increased MVPA accuracy) for the trained orientation in superficial rather than middle or deeper layers in V1 (Figure 4, Figure 5A). In particular, a three-way repeated measures ANOVA (orientation × session × layer) on the MVPA accuracy showed a significant three-way interaction (F(2,24) = 4.244, *p* = 0.026). Two-way repeated measures ANOVAs (orientation × session) showed a significant interaction in superficial V1 layers (F(1,12) = 12.223, *p* = 0.004), but not in middle (F(1,12) = 0.012, *p* = 0.913), nor deeper (F(1,12) = 0.446, *p* = 0.517) layers. Further, we observed enhanced discriminability (i.e. MVPA accuracy) for the trained orientation in superficial (t(12) = −2.665, *p* = 0.021), but not middle (t(12) = −0.783, *p* = 0.449), nor deeper (t(12) = −0.489, *p* = 0.633) layers. In contrast, we did not observe any significant learning-dependent changes for orientations presented in the untrained location. In particular, there was no significant three-way interaction (orientation × session × layer, F(2,24) = 0.603, *p* = 0.555) nor any significant orientation × session interactions across V1 layers (superficial layers: F(1,12) = 0.053, *p* = 0.821; middle layers: F(1,12) = 0.538, *p* = 0.478; deeper layers: F(1,12) = 2.211, *p* = 0.163). Taken together, our results showed that learning-dependent changes in orientation representations in superficial V1 layers were specific to the trained orientation and location. It is unlikely that these learning-dependent changes in visual representations were due to differences in attention related to task difficulty before vs. after training, as participant performance was matched across scanning sessions.

**Figure 4.**
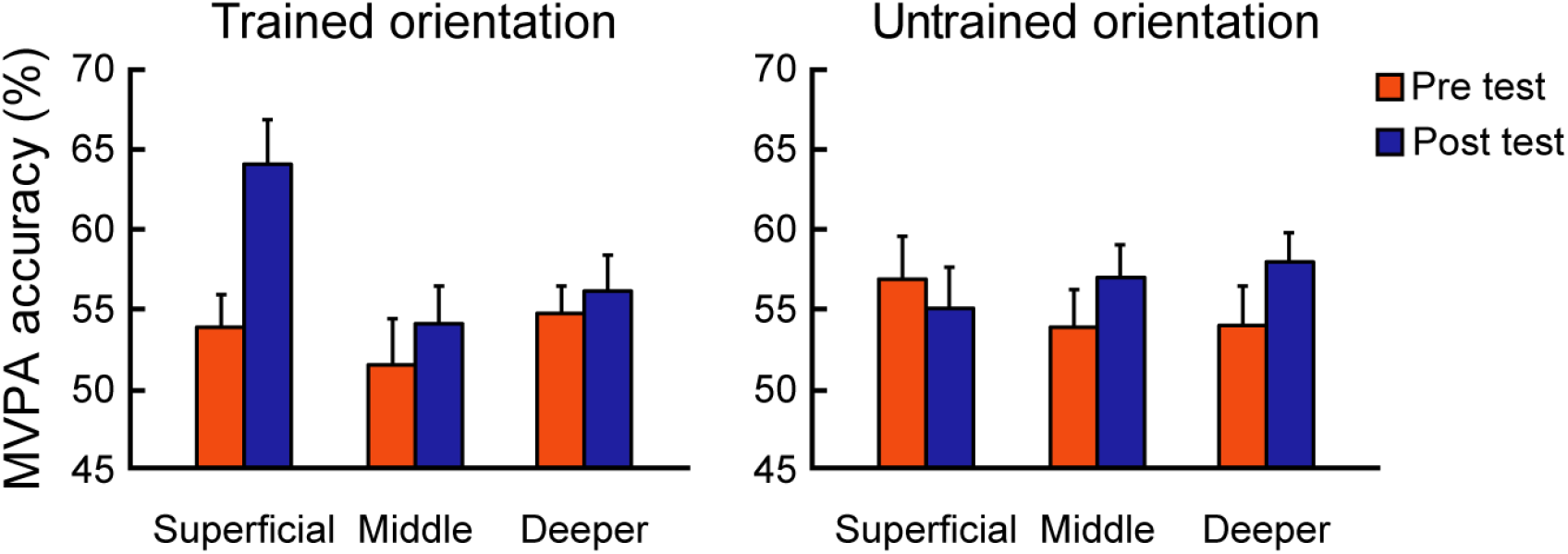
MVPA results before and after training across V1 layers. MVPA accuracy across V1 layers for the trained and untrained orientations presented at the trained location. To further validate our classification results, we trained the classifier after shuffling the labels of the training data set for 5000 times. This analysis returned classification accuracies that did not differ significantly from chance (all *ps* > 0.618, FDR corrected), suggesting that our MVPA analysis extracted reliable voxel pattern information from the ROIs tested. Error bars indicate standard error of the mean across participants.

**Figure 5.**
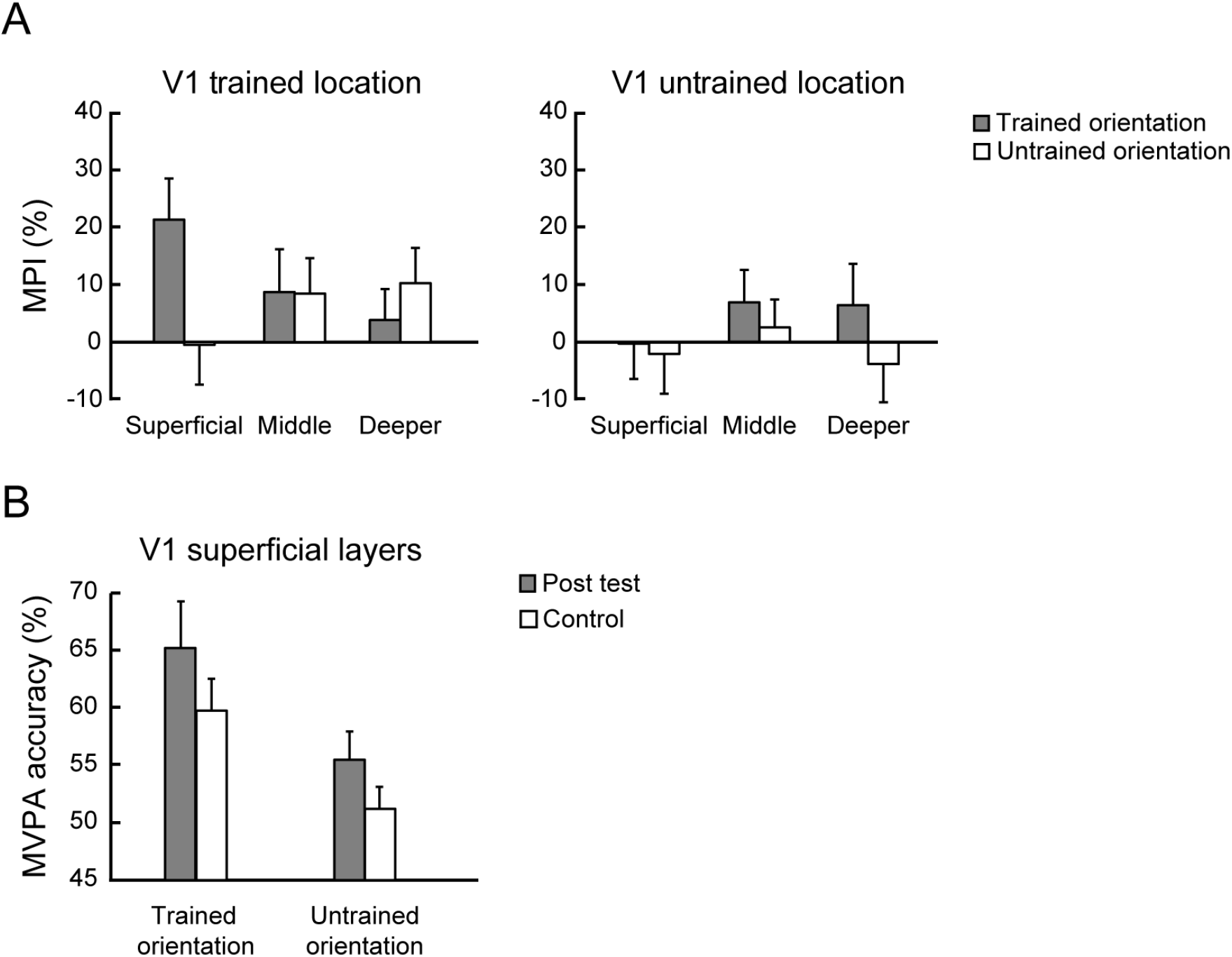
Learning-dependent changes in V1. (A) Mean improvement index (MPI: (post-test accuracy – pre-test accuracy) / pre-test accuracy × 100%) for the trained and untrained orientations across V1 layers. The left panel shows significantly higher MPI for the trained than the untrained orientation at the trained location in superficial V1 layers, as indicated by significantly higher MPI for the trained than untrained orientation in the superficial layers (t(12) = 3.218, *p* = 0.007). The right panel shows no significant differences in MPI across layers for orientations presented at the untrained location. In particular, there was no significant differences in MPI for trained vs. untrained orientations in the superficial layers (t(12) = 0.396, *p* = 0.699). (B) MVPA accuracy for the trained and untrained orientation at the trained location after training (post-test) compared to the control experiment in superficial layers of V1. Error bars indicate standard error of the mean across participants.

### Complementary and control analyses

To further validate our results and control for potential confounds we conducted the following additional analyses.

First, it has been shown that the overall BOLD signal as measured by GE-EPI is higher at the cortical surface due to vascular contributions^32^ resulting in loss of spatial specificity^33^. Here, we combined several approaches to reduce this superficial bias by removing voxels with low temporal signal to noise ratio and high t-statistic for stimulation contrast (*see Methods, correcting for vascular effects for details*). We then z-scored each voxel’s time course to account for possible differences in signal strength and variance due to thermal or physiological noise across layers while preserving differences between conditions^29^. These corrections resulted in similar BOLD magnitude and multi-voxel pattern classification accuracy before training across layers (Figure 3E), suggesting that our approach for correcting vasculature-related effects controlled substantially for the superficial bias. In particular, consistent with previous studies^34^ showing reduced superficial bias for MVPA classification measures, we did not observe any significant differences in MVPA accuracy between trained and untrained orientations before training (e.g. orientation × location × layer interaction: F(2,24) = 0.891, *p* = 0.423; main effect of layer: F(2,24) = 0.287, *p* = 0.753). Thus, it is unlikely that our MVPA results after vasculature correction were significantly confounded by the superficial bias.

Second, we applied a spatial regression approach^35,36^ to control for signal contribution from draining veins. In particular, intra-cortical veins running perpendicular to the cortical surface are known to drain blood from deeper layers of the cortex to larger pial veins situated along the grey matter surface, resulting in loss of spatial specificity and intra-layer BOLD signal contamination. To unmix the signal from adjacent layers, for each voxel in the superficial layers, we found the nearest neighbor in the middle layer. We regressed out the mean time course of these voxels assigned to middle layers from the time course of voxels assigned to superficial layers. MVPA analysis following this correction showed a significant interaction between orientation and session (F(1,12) = 14.357, *p* = 0.003, Figure S2A), consistent with learning-dependent changes in superficial layers. Further, we observed enhanced discriminability in superficial layers after correction for the trained orientation (t(12) = −2.292, *p* = 0.041), but not the untrained orientation (t(12) = 0.246, *p* = 0.810). Learning-dependent changes in superficial V1 layers remained significant after these corrections, suggesting that our results are unlikely to be significantly confounded by vasculature-related artifacts.

Third, comparing mean normalized fMRI responses (Figure S3) across orientations and sessions did not show any significant results (three-way interaction (orientation × session × layer): F(4,48) = 1.259, *p* = 0.299, two-way interaction (orientation × session) in the superficial layers: F(2,24) = 0.814, *p* = 0.455, middle layers: F(2,24) = 0.934, *p* = 0.407 and deeper layers: F(2,24) = 0.389, *p* = 0.682), suggesting that the learning-dependent effects we observed reflect changes in orientation-specific representations across voxel patterns rather than mean univariate fMRI responses. Fourth, we corroborated our results using a correlation-based pattern analysis^37^ that showed learning-dependent changes in superficial V1 layers for the trained compared to the untrained orientation (Figure S2B). Specifically, Fisher z comparisons showed a significant orientation × session interaction in superficial V1 layers (F(1,12) = 6.069, *p* = 0.030), but not in middle (F(1,12) = 2.382, *p* = 0.149) nor deeper layers (F(1,12) = 1.227, *p* = 0.290).

Taken together, these results demonstrate enhanced orientation-specific representations in superficial rather than middle or deeper V1 layers after training, suggesting that learning alters read-out rather than input processing in V1. Similar learning-dependent effects with stronger learning-dependent changes in superficial layers were observed across visual areas (V1, V2, V3, V4, Figure S4). In particular, a repeated measures ANOVA showed no significant ROI × orientation × session × layer interaction (F(2,24) = 1.459, *p* = 0.252), but a significant orientation × session × layer interaction (F(2,24) = 5.305, *p* = 0.012), suggesting similar orientation-specific learning effects in superficial layers across visual areas.

### Learning-dependent changes independent of task-context in the visual cortex

Our results showed learning-dependent changes in orientation-specific representations when participants performed a fine orientation discrimination task. To test whether the task performed by the participants is critical for this orientation-specific plasticity, we tested a subset of participants (n = 8) in a second post-training fMRI session while performing a control task (i.e. contrast change detection task) on identical stimuli to those presented to the participants during the post-training fMRI session. Before this additional scanning session, we conducted behavioral tests to ensure that perceptual improvement was retained. We observed that threshold performance (Mean = 1.78°, SD = 0.43°) did not differ significantly from the mean threshold of training day 5 (Mean = 1.69°, SD = 0.37°, paired t-test, t(7) = −1.553, *p* = 0.164). Further, we matched task difficulty in the contrast change detection task (~79.4%) to performance in the orientation discrimination task during the post-training fMRI session to ensure that the two post-training scanning sessions did not differ in task difficulty.

MVPA analysis across cortical layers in V1 showed that the learning-dependent changes we observed in orientation-specific representations in superficial layers were maintained when participants performed the control task. In particular, a two-way repeated measures ANOVA (orientation × task) on the MVPA accuracy showed a significant main effect of orientation (F(1,7) = 19.140, *p* = 0.003) in superficial V1 layers (Figure 5B). Neither the main effect of task (F(1,7) = 2.608, *p* = 0.150) nor the interaction effect (F(1,7) = 0.082, *p* = 0.783) were significant. We did not find any significant main or interaction effects in the deeper or middle layers of V1 (all *ps >* 0.071). These results suggest that learning-dependent changes in orientation-specific representations in superficial V1 layers are independent of task context, consistent with previous neurophysiology results^9^ showing learning-dependent changes in neural tuning in primary visual cortex during a fixation task.

### Learning-dependent changes in intraparietal cortex

We next considered learning-dependent changes in decision-making related areas^6,38^ outside the visual cortex. In particular, we focused on IPS1 and IPS2 that have been implicated in perceptual decision making^38^. Using MVPA, we tested for learning-dependent changes in orientation representations across cortical depth. Figure 6 shows learning-dependent changes for the trained compared to the untrained orientation (i.e. increased MVPA accuracy for the trained orientation in the trained location) in middle rather than superficial or deeper layers. In particular, we observed a significant orientation × session interaction in middle layers of IPS (F(1,12) = 6.324, *p* = 0.027), but not in superficial (F(1,12) = 0.502, *p* = 0.492), nor deeper (F(1,12) = 2.452, *p* = 0.143) layers. Further, we observed significantly increased MVPA accuracy after training for the trained orientation in the middle (t(12) = −3.432, *p* = 0.005), but not superficial (t(12) = 0.392, *p* = 0.702), nor deeper (t(12) = −1.137, *p* = 0.278) layers, suggesting enhanced discriminability for the trained orientation in middle layers. In contrast, we did not observe any significant learning-dependent changes for the untrained location. In particular, there was no significant orientation × session interactions across IPS layers (superficial layers: F(1,12) = 0.136, *p* = 0.718; middle layers: F(1,12) = 1.053, *p* = 0.325; deeper layers: F(1,12) = 1.628, *p* = 0.226). Comparing learning-dependent changes in visual and posterior parietal cortex showed dissociable results. That is, we observed learning-dependent changes in superficial layers of visual cortex, while middle layers of posterior parietal cortex, as indicated by a significant ROI (V1, IPS) × session × layer interaction for the trained orientation (repeated measures ANOVA, F(2,24) = 5.872, *p* = 0.008), but not the untrained orientation (repeated measures ANOVA, F(2,24) = 1.933, *p* = 0.167).

**Figure 6.**
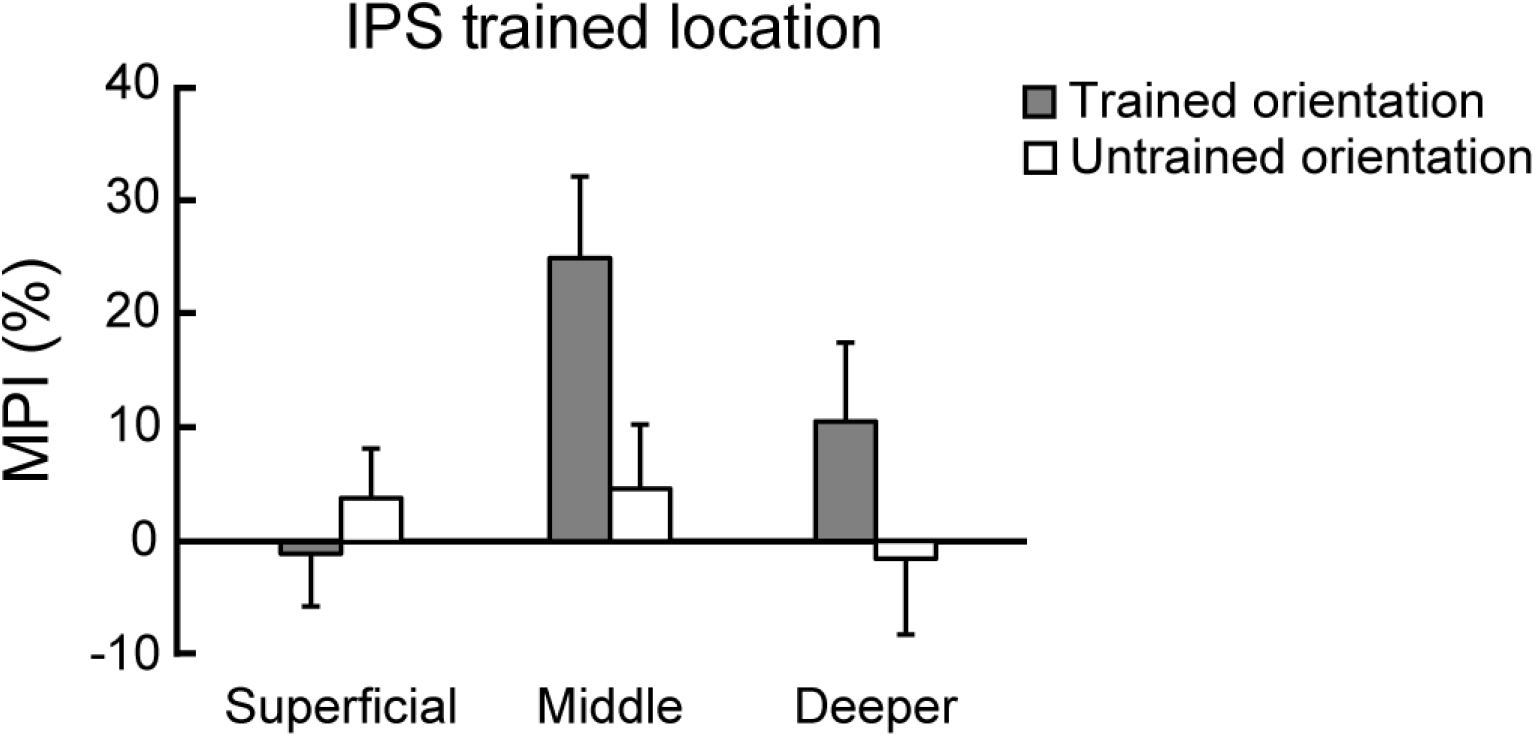
Learning-dependent changes in IPS. Mean improvement index across IPS layers for the trained and untrained orientations presented at the trained location. The results showed significantly higher MPI for the trained than the untrained orientation at the trained location in the middle IPS layers (t(12) = 2.861, *p* = 0.014), but not in superficial (t(12) = − 0.695, *p* = 0.500), nor deeper (t(12) = 1.382, *p* = 0.192) layers. Error bars indicate standard error of the mean across participants.

### Informational connectivity analysis

Our results so far showed learning-dependent changes in orientation-specific representations in the superficial layers of V1 and middle layers of IPS, suggesting that learning modifies read-out rather than input signals in visual cortex, while input signals in posterior parietal cortex. Based on these results, we asked whether learning enhances functional connectivity between visual and posterior parietal cortex. To test this hypothesis, we employed an informational connectivity analysis^36^ and tested whether V1 and IPS shared synchronous discriminability of multi-voxel patterns that changed with training. Consistent with previous studies^19,20,26^, we contrasted two possible functional connectivity mechanisms: a) feedforward learning, as indicated by changes in connectivity between superficial V1 layers and middle IPS layers, b) feedback learning, as indicated by changes in connectivity between V1 deeper layers and IPS deeper layers. We did not test functional connectivity between V1 superficial layers and deeper layers of higher areas, as it is known to relate to both feedback and feedforward processing (Figure 1).

Following previous studies employing an informational connectivity analysis^36^, we interrogated the MVPA classifiers and extracted the distance from the hyperplane for the mean pattern signal per block. For each layer per ROI we generated a time course of distance values across blocks, regressed out the distance from other layers within the ROI and calculated the partial Spearman correlation between V1 and IPS layers across blocks (Figure 7A). Our results showed enhanced feedforward compared to feedback connectivity between V1 and IPS after training. A repeated measures ANOVA (Fisher’s z) showed a significant pathway (feedforward, feedback) × orientation (trained vs. untrained) × session (pre-test, post-test) interaction (F(1,12) = 8.912, *p* = 0.011, Figure 7B).

**Figure 7.**
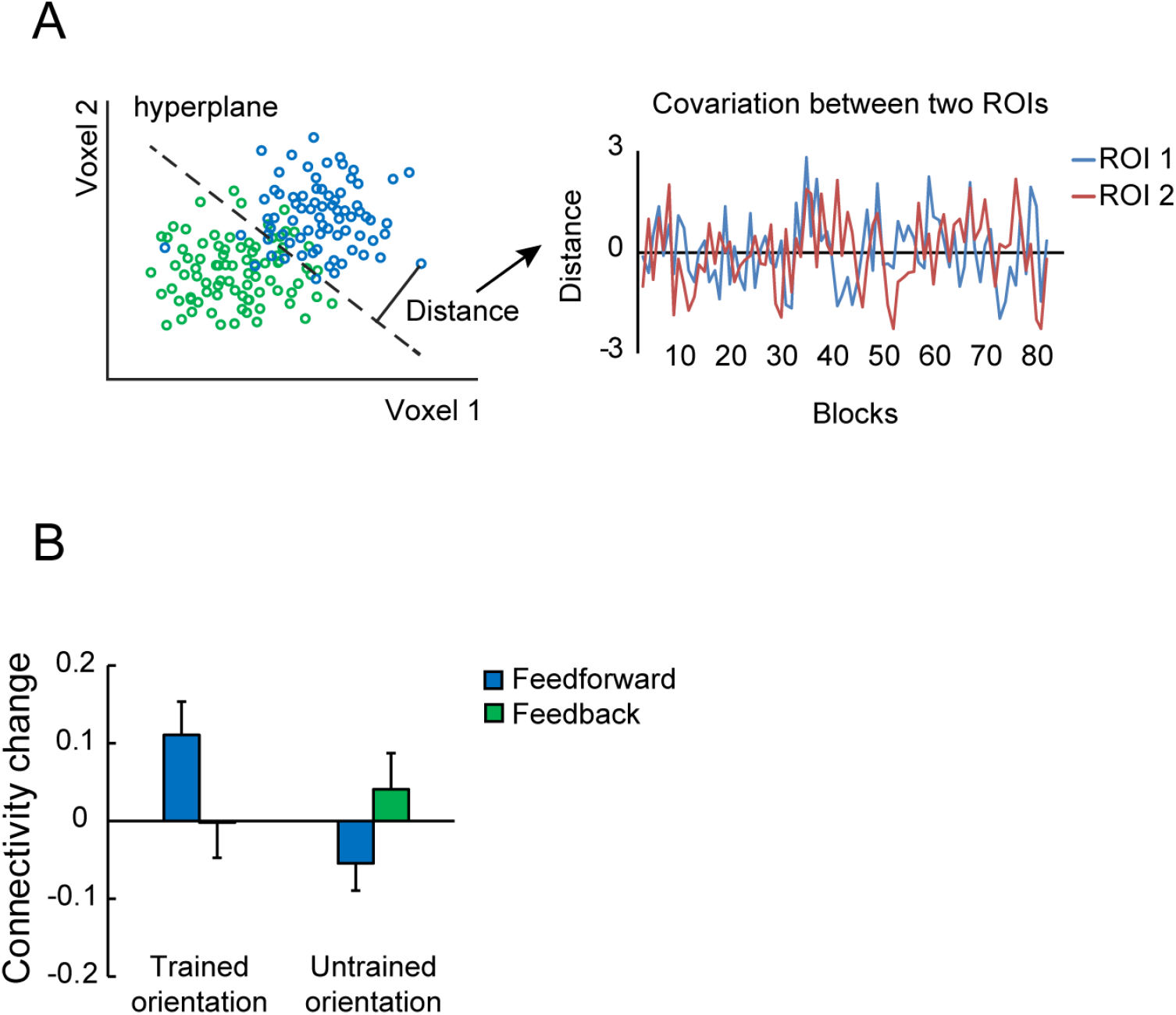
Informational connectivity analysis. (A) Schematic illustration of the procedure followed for the MVPA-based functional connectivity analysis. For each ROI and block, the distance to the hyperplane was used as an index of pattern discriminability (left panel). Spearman correlation was used to calculate covariance between two ROIs (right panel). (B) Learning-dependent changes (Fisher’s z post-minus pre-test) in functional connectivity between superficial V1 layers and middle IPS layers (feedforward connectivity) and between deeper V1 and IPS layers (feedback connectivity) for the trained and untrained orientations. Error bars indicate standard error of the mean across participants.

In particular, we observed enhanced feedforward connectivity between superficial V1 layers and middle IPS layers after training. A repeated measures ANOVA (Fisher’s z) showed a significant orientation × session interaction (F(1,12) = 5.771, *p* = 0.033), but no significant main effect of orientation (F(1,12) = 1.218, *p* = 0.291) nor session (F(1,12) = 2.326, *p* = 0.153). Post-hoc comparisons showed enhanced connectivity for the trained orientation (t(12) = −2.599, *p* = 0.023), but not the untrained orientation (t(12) = 1.560, *p* = 0.145). In contrast, we did not observe any significant learning-dependent changes in feedback connectivity between deeper V1 and deeper IPS layers (orientation × session interaction: F(1,12) = 0.587, *p* = 0.458, main effect of session: F(1,12) = 0.299, *p* = 0.595, main effect of orientation: F(1,12) = 1.223, *p* = 0.290). This enhanced connectivity between superficial layers of V1 and middle layers of IPS after training suggests that training alters feedforward information processing from sensory to decision-related areas to support learning-dependent improvement in the fine orientation discrimination task.

## Discussion

Despite the key role of learning in optimizing perceptual decisions, an ability known as perceptual learning, we lack a mechanistic account of how training molds the adult human brain. Evidence on experience-dependent plasticity mechanisms remains controversial with some studies suggesting that learning modifies sensory encoding, while others proposing top-down influences via feedback^5,39,40^. Conventional brain imaging techniques have been unable to differentiate these possibilities due to limited spatial resolution^18^. Here, we capitalize on the sub-millimeter resolution of 7T laminar fMRI to interrogate plasticity mechanisms across cortical depths that are known to be associated with dissociable neural computations. Our results provide evidence for recurrent experience-dependent plasticity mechanisms, offering an alternate proposal for the neural instantiation of perceptual learning, by contrast to the long-standing debate on encoding vs. feedback mechanisms of sensory plasticity.

Combining 7T laminar fMRI with MVPA we demonstrate enhanced decoding of orientation-specific representations in superficial rather than middle layers of V1, suggesting that learning modifies read-out rather than input signals in visual cortex^26^. Previous fMRI studies using multi-voxel pattern classification approaches have shown that learning fine feature discriminations increases the discriminability of neural representations^12,41^. Other brain imaging studies^11,13,42,43^ have shown learning-dependent changes in overall BOLD signal in V1. These fMRI results have been interpreted broadly in support of an early neural locus of perceptual learning. However, we still lack a mechanistic account of experience-dependent plasticity in the human brain, as fMRI at standard resolution does not allow us to discern encoding from read-out processes. Our layer-specific fMRI results propose that learning alters the processing of read-out signals in superficial V1 layers rather than stimulus encoding in middle layers. It is unlikely that this learning-dependent plasticity in superficial layers relates to stronger orientation selectivity in superficial V1 layers, as recent quantitative measurements show nearly uniform distribution of orientation selectivity across V1 layers than reported in early physiological studies^44^. Further, we found that MVPA accuracy did not differ significantly across layers before training, suggesting that learning-dependent changes in superficial V1 layers could not be due to differences in orientation selectivity across V1 layers.

A possible explanation is that learning-dependent changes in superficial layers reflect top-down influences via feedback^45,46^. It is possible that neurons with cell bodies in deeper layers and dendrites projecting to the superficial layers result in learning-dependent changes in BOLD signals in superficial layers^20,21^. However, this interpretation is less likely for three main reasons. First, we did not observe learning-dependent changes in deeper V1 layers that are known to receive long-range feedback. Second, our functional connectivity analysis showed enhanced feedforward rather than feedback connectivity in occipito-parietal circuits. Third, consistent with a previous physiological study^9^, learning-dependent changes in superficial V1 layers were maintained independent of task context; that is, these changes were evident not only when participants performed the orientation discrimination task, but also when they performed a contrast detection task (i.e. control experiment) that does not involve task-related feedback on the trained stimulus dimension (i.e. orientation). It is possible that top-down influences, via feedback connections to superficial layers, play a role in shaping visual feature templates at earlier than later stages of training on the discrimination task^47^. However, interrogating the representations of trained features after training reveals learning-dependent changes in superficial layers that are independent of task context.

An alternate proposal is that orientation-specific learning-dependent changes in superficial V1 layers are due to recurrent processing related to iso-orientation inhibition^48^; that is, suppression of neurons that are selective for the same orientation across columns. Iso-orientation inhibition is shown to be more pronounced in superficial layers and support orientation tuning via horizontal connections between V1 columns^19,23,24^. It is possible that iso-orientation suppression supports enhanced neural tuning to the trained orientation in superficial V1 layers, suggesting a recurrent experience-dependent plasticity mechanism that alters orientation-specific representations within visual cortex. These results are consistent with computational models proposing that training sharpens neural tuning by altering recurrent connections (i.e. reducing excitatory or increasing inhibitory connections) near the trained orientation^25,49^ and neurophysiological studies showing changes in orientation tuning due to training in visual cortex ^9,10^. Further horizontal connections in superficial V1 layers have been suggested to support recurrent processing in the context of figure-ground segmentation^26^ and experience-dependent plasticity in the context of contour integration^2^. In particular, boundary detection was evidenced in superficial layers ~30 ms after the initial visual response in the middle layers^26^. Future studies combining laminar fMRI with EEG in humans have the potential to test whether learning-dependent changes in superficial layers follow early sensory processing in middle layers, revealing the dynamics of recurrent experience-dependent plasticity^26,27^.

It is important to note that despite the advances afforded by laminar fMRI, GE-EPI is limited by vasculature contribution to BOLD signals at the cortical surface resulting in loss of spatial specificity^33^. Here, we demonstrate layer-specific learning-dependent changes following several control analyses for these potential confounds, suggesting that our results are unlikely to be confounded by vasculature-related artefacts. Our results are consistent with previous laminar imaging studies showing BOLD effects in superficial layers in a range of tasks^28,29,45,46^ and could not be simply attributed to differences in attention due to task difficulty, as participant performance was matched across sessions (pre vs. post-training). Future work could exploit recent advances in CBV imaging using vascular space occupancy (VASO)^50^ to enhance the spatial specificity of laminar imaging in the human brain.

Extending beyond the visual cortex, we demonstrate learning-dependent changes in posterior parietal cortex regions (IPS) that have been suggested to play a key role in perceptual decision making^6^. In particular, we observed learning-dependent changes in fMRI activation patterns in middle layers of IPS regions, suggesting that training alters input signals to posterior parietal cortex. These results are consistent with previous neurophysiological^6^ and human brain imaging studies^38^ that show learning-dependent changes in intraparietal cortex for perceptual decision-making. Our findings provide new insights in understanding the finer scale circuit that mediates experience-dependent plasticity across human brain systems, involving both sensory and decision-related areas^51^.

Further, we interrogated learning-dependent changes in functional connectivity^38,52^. We tested whether training alters the functional connectivity within this circuit at the finer scale of layer-to-layer interactions. Computational approaches have proposed that training strengthens the connections between the most informative neurons in sensory areas and decision-related areas via Hebbian learning, resulting in re-weighting of sensory signals in visual cortex^5^. In contrast, the reverse hierarchy theory proposes that learning is implemented by top-down influences to visual processing via long-range feedback from downstream areas^17^. To test these hypotheses, we interrogated the layer-to-layer functional connectivity between visual and posterior parietal cortex that allows us to compare feedforward vs. feedback processing based on known anatomical connectivity models^19,20,53^. We demonstrate that learning strengthens feedforward connectivity between superficial V1 layers and middle layers of IPS, consistent with previous studies showing that ascending projections of V1 originate predominantly from the superficial layers and ascending projections to IPS mainly terminate in middle layers^19,20,26^. In contrast, we did not find any significant changes in feedback connectivity between deeper V1 and deeper IPS layers, consistent with the lack of significant learning-dependent changes in deeper V1 layers. Taken together, these results suggest that learning fine feature differences is implemented by re-weighting mechanisms of visual plasticity rather than long-range feedback from decision-related to visual areas. Corroborating evidence comes from the results of our control experiment showing that learning-dependent changes in superficial V1 layers were maintained when observers performed a contrast change detection task that does not involve orientation judgments and therefore does not engage decision-related feedback on the trained stimulus dimension.

In sum, combining ultra-high field 7T imaging with a classic perceptual learning paradigm, we provide evidence for recurrent mechanisms of experience-dependent plasticity that gate perceptual decision making in the adult human brain. These mechanisms support re-weighting of input signals in the visual cortex that are read-out by posterior parietal cortex to inform improved perceptual decisions due to training. Interrogating experience-dependent plasticity at the finer resolution afforded by UHF imaging using training paradigms that have been extensively tested in both animals and humans, we provide the first insights in bridging the gap between animal studies of micro-circuit plasticity and human fMRI studies of macro-scale network re-organization. Understanding the mechanisms of experience-dependent plasticity in the adult brain across scales and species is critical for designing effective training interventions that support lifelong learning and adaptive behavior.

## Materials and Methods

### Participants

Fifteen participants (7 females; mean age: 27.87 years and SD: 3.83 years) took part in this study. Data from two participants were excluded from further analysis due to excessive head movement (See MRI data analysis) and technical issues during acquisition. As there are no previous UHF imaging studies on perceptual learning, we determined the sample size based on a previous 3T fMRI study on perceptual learning using an orientation discrimination task (N = 12)^12^. All participants had normal or corrected-to-normal vision, and were right-handed. Participants were naïve to the aim of the study, gave written informed consent and received payment for their participation. The study was approved by the University of Cambridge Ethics committee.

### Stimuli and Apparatus

Stimuli comprised oriented sinusoidal gratings that were presented at an eccentricity of 5°, in the left or the right visual field against a uniform gray background. Gratings of random phase had a fixed diameter of 6°, contrast of 0.8, spatial frequency of 1 cycle/degree. The contrast decreased to zero over the outer 0.5° radius of the gratings.

Experiments were controlled using MATLAB and Psychophysics toolbox 3.0^54,55^. For the behavioral sessions, stimuli were presented on a 21-inch CRT monitor (1600 × 1200 pixel resolution, 85 Hz frame rate) at a distance of 110 cm. Gamma correction was applied to the monitor. For the fMRI scans, stimuli were presented using a projector and a mirror setup (1920 × 1080 pixel resolution, 100 Hz frame rate) at a viewing distance of 110 cm. Angular stimulus size was the same across behavioral and fMRI sessions.

### Experimental design

The study comprised a pre-test (2 sessions, 1 behavioral test, 1 fMRI test), a training (5 sessions), a post-test (2 sessions, 1 behavioral test, 1 fMRI test) and a control (2 sessions: 1 training, 1 fMRI test) phase (Figure 2A). All training and post-test sessions were conducted on consecutive days.

We employed a two-interval forced choice (2IFC) orientation discrimination task (Figure 2B). Each trial began with a fixation cross for 200 ms followed by the sample and test gratings that were presented sequentially for 200 ms each and separated by a 600 ms inter-stimulus interval (ISI). Participants were asked to fixate and report (by key press) within 1270 ms after the onset of the test grating whether it was tilted clockwise or counter-clockwise relative to the sample stimulus.

Participants’ performance in the task was measured using a 3-down-1-up staircase with 15 reversals converging at 79.4% performance. The reference orientation for the trained and untrained stimuli was 55° or 125°. We added a uniformly distributed random jitter within ±5° to the reference orientation across trials to ensure that participants compared two gratings in each trial, rather than the test grating to a fixed reference orientation. The training reference orientation (55° vs. 125°) was counterbalanced across participants. Participants were tested with a control orientation (0°, vertical) that differed equally from the trained and untrained orientation (55° or 125°). This allowed us to test orientation-specific pattern changes in fMRI signals due to training (i.e. learning-dependent changes to the trained vs. untrained orientation), by comparing separately the trained vs. the untrained orientations to the control orientation.

#### Behavioral Tests

To familiarize participants with the task before testing, each participant performed a 30-trial practice run (5 trials per condition, i.e. three different reference orientations at two different locations) using a fixed above-threshold angle difference (8°). For both the pre- and post-training test, participants performed the orientation discrimination task for 12 test staircase runs (2 runs per condition in random order). For each condition, the starting angle difference between sample and test stimulus for the first run was 5°. For the second run, the starting angle difference was determined by the threshold in the preceding run. The discrimination threshold for each condition was the mean threshold across two runs. No feedback was provided to the participants during the test phase.

#### Behavioral Training (5 sessions)

We trained participants on the orientation discrimination task (16 staircases per session, ~1 h) with gratings presented at the same orientation and location throughout training. The starting angle difference between sample and test stimuli for the first staircase of the first training session was 5°. For the remaining staircases, the starting angle difference was determined by the threshold of the preceding staircase. Training location (i.e., left vs. right visual field) and orientation (i.e., 55° vs. 125°) were counterbalanced across participants. Participants were given auditory error feedback per trial.

#### fMRI sessions

Before and after training in the lab, participants completed 8-10 runs of the orientation discrimination task during scanning. For each participant, we also collected data from an anatomical scan and a retinotopic mapping scan.

For the orientation discrimination task, each run started with a fixation block (12.36 sec) followed by one block for each of the six conditions and a fixation block. This sequence of fixation and condition blocks was repeated four times in each run. Each condition block lasted 12.36 sec and comprised gratings presented at the trained, untrained or control orientation at one of two locations (trained vs. untrained location). The order of orientations was randomized across the six condition blocks and the stimulus location alternated between blocks. For each block, participants completed five trials of the orientation discrimination task. The task parameters (i.e. sample and test duration) were the same as for the behavioral tests and no feedback was provided to the participants. The fixed angle difference between sample and test stimuli for each condition was determined by the preceding behavioral session. This allowed us to match task difficulty (~79.4%) before and after training.

#### Control experiment

To test the task specificity of the learning effect, eight participants completed an additional training and fMRI session on consecutive days to ensure that learning was maintained during a control task. The procedure for this control-task session was identical to that for the post-training scan, with the exception that participants performed a contrast change detection task. Participants were required to press a key within 1000 ms of detecting a contrast change on the stimuli. The magnitude of contrast change was estimated for each participant during the anatomical scan to ensure similar task difficulty (~79.4%) between the orientation discrimination task and the contrast detection task. The estimated magnitude was fixed and used throughout the control-task fMRI session.

### MRI data acquisition

Imaging data were acquired at the Wolfson Brain Imaging Centre, University of Cambridge, on a Siemens 7T Terra scanner with a 32-channel phased-array head coil (Nova Medical, Inc., Wilmington, MA, USA). For each participant, anatomical images were acquired using MP2RAGE T1-weighted sequence (TR = 5000 ms, TE = 2.56 ms, FOV = 208 × 208 mm^2^, resolution 0.65 × 0.65 × 0.65 mm^3^, number of slices: 240, slice orientation: sagittal). Functional scans were acquired using a 2D Gradient Echo, Echo Planar Imaging (GE-EPI) sequence^56^ (TR = 2060 ms, TE = 26.4 ms, FOV = 148 × 148 mm^2^, flip angle: 70°, resolution 0.8 × 0.8 × 0.8 mm^3^, number of slices: 56, partial Fourier = 6/8, GRAPPA factor = 3, Multi-Band factor = 2, bandwidth = 1034 Hz/Pixel, echo spacing = 1.09 ms). The field of view covered occipito-temporal and posterior parietal areas; manual shimming was performed prior to the acquisition of the functional scans.

### Behavioral data analysis

Performance was measured by the 3-down-1-up staircase with 15 reversals. The mean angle difference of the last 8 reversals was taken as the threshold of each staircase run. The measured orientation discrimination thresholds were used as the dependent factor. Using a within-subject factorial design, we manipulated three independent factors, the reference orientation (trained and untrained orientation), stimulus location (trained and untrained location) and test session (pre-test, post-test), to evaluate the learning effect and learning specificity. Further, we calculated the mean percent improvement index (MPI, (pre-test threshold – post-test threshold) / pre-test threshold × 100%)) for each condition. For statistical analysis, we used repeated measures ANOVAs to compare across conditions.

### MRI data analysis

#### Anatomical data analyses

T1-weighted anatomical data was used for coregistration and 3D cortex reconstruction. Grey and white matter segmentation was obtained on the MP2RAGE images using FreeSurfer (http://surfer.nmr.mgh.harvard.edu/) and manually improved for the regions of interest (i.e., V1, V2, V3, V4, and IPS) using ITK-SNAP (www.itksnap.org). The refined segmentation was used to obtain a measurement of cortical thickness. Following previous studies, we assigned voxels to three layers (superficial, middle, deeper) using the equi-volume approach^57,58^ as implemented in BrainVoyager (Brain Innovation, Maastricht, The Netherlands). This approach has been shown to reduce misclassification of voxels to layers, in particular for regions of interest presenting high curvature. Information from the cortical thickness map and gradient curvature was used to generate four grids at different cortical depths (ranging from 0: white matter, to 1: grey matter). Mapping of each voxel to a layer was obtained by computing the Euclidean distance of each grey matter voxel to the grids: the two closest grids represent the borders of the layer to which a voxel is assigned (Figure 3B). The anatomical image was aligned to the functional data using the boundary-based registration^59^. We assessed the alignment and manually corrected if necessary.

#### Functional data analyses

The GE-EPI functional data were analysed using BrainVoyager (version 20.6, Brain Innovation, Maastricht, The Netherlands) and custom MATLAB (The MATHWORKS Inc., Natick, MA, USA) code. The first two volumes at the beginning of each run were discarded to ensure that the longitudinal magnetization reached steady state. The functional data were corrected for distortions due to non-zero off-resonance field (at the beginning of each functional run, five volumes with inverted phase encoding direction were acquired and used to estimate a voxel displacement map that was subsequently applied to the functional data using COPE (Correction based on Opposite Phase Encoding, BrainVoyager, Brain Innovation). The distortion-corrected data underwent slice-timing correction, head motion correction (the single band image acquired at the beginning of the first run was used as the reference in the alignment), high-pass temporal filtering (using a GLM with Fourier basis set at 2 cycles) and removal of linear trends. We then aligned the functional data across sessions. To validate the alignment, we calculated the mean EPI image of each functional run for each region of interest (ROI) and estimated the spatial correlation between these mean EPI images. We performed manual adjustment of the alignment if the spatial correlation was below 0.85 and excluded data from one participant for whom the alignment could not be improved manually.

#### Regions of Interest definition

We used the data from the retinotopic mapping scan to identify visual areas based on standard phase-encoding methods. Participants viewed rotating wedges that created travelling waves of neural activity^60,61^. Due to limited coverage during acquisition, area V4 was identified for 8 of the 13 participants included in the analysis. Thus, for further analyses we combined the data from V2, V3 and V4 for each individual participant. Further, we defined regions in the intraparietal sulcus (IPS1, IPS2) given their functional relevance for perceptual learning ^6^. Intraparietal regions (IPS1 and IPS2) were defined for each participant based on anatomical templates provided by Benson (https://hub.docker.com/r/nben/occipital_atlas/)^62^. This procedure uses the individual participant-based segmentation obtained with FreeSurfer and an anatomical probabilistic template, to estimate the best location for the region of interest (i.e. IPS). Each IPS subregion was subsequently inspected to ensure consistent definition across participants.

For each of the visual cortex ROIs, we modelled BOLD signals using a GLM with two regressors (i.e., left vs. right visual field) and included the estimated head motion parameters as nuisance regressors. The resulting t-statistical map was thresholded (t = 2.58, *p* = 0.01) to select voxels within each ROI that responded strongly to the lateralized stimulus presentation, consistent with location specificity in visual cortex. For IPS, we selected voxels that responded to the task irrespective of stimulus location (i.e. task vs. fixation, t = 1.64, *p* = 0.10).

#### Correcting for vasculature-related effects

Voxel selection within each ROI was further refined by excluding voxels that were confounded by vasculature effects that are known to contribute to a superficial bias in the measured BOLD signal; that is, increased BOLD with increasing distance from white matter. In particular, it has been shown that the BOLD signal measured using GE-EPI (i.e. T2* weighted) is confounded by macro- and micro-vasculature signals^32,63,64^.

The macro-vasculature contribution is due to veins penetrating the grey matter and running through its thickness, as well as large pial veins situated along the surface of the grey matter^65^. This results in increased sensitivity (i.e., strong BOLD effect) but decreased spatial specificity of the measured signal. The latter can be understood by the mechanics of the draining veins carrying deoxygenated haemoglobin downstream from the true neuronal site of neural activation, leading to a response spatially biased towards the pial surface, an effect known as superficial bias. Here, we took the following approach to reduce superficial bias due to vasculature contributions. First, following previous work^66^, we computed the temporal signal to noise ratio (tSNR) for each voxel in each ROI (V1 V2, V3, V4 and IPS respectively). We used tSNR to identify voxels near large veins that are expected to have large variance and low intensity signal due to the local concentration of deoxygenated haemoglobin resulting in a short T2* decay time (i.e., dark intensity in a T2* weighted image). We identified voxels with low tSNR, checked their correspondence with voxels of lower intensities on the T2* weighted images. Second, it has been shown that high t-values on a fMRI statistical map are likely to arise from large pial veins^67,68^. Therefore, voxels with low tSNR values or t-score values above the 90^th^ percentile of the t-score distribution obtained by the GLM described above were removed from further analysis. We used these two approaches to correct the BOLD signal from confounding vasculature effects.

#### Univariate analysis

For each participant, test session, run and condition, we extracted the z-scored fMRI responses between the 4^th^ and 8^th^ TR (i.e. 6.18 – 14.42 s) after block onset. This time window captured the peak of the hemodynamic responses to the visual stimuli. The normalized fMRI responses were averaged across time points, blocks and runs for each condition and each session. Repeated-measures ANOVA was used to test the univariate difference across conditions.

#### Multivariate pattern analysis

We used multivariate pattern analysis (MVPA) to decode: a) trained vs. control orientation, b) untrained vs. control orientation. For each ROI and participant, we calculated per voxel a t-score statistic by comparing activity for stimuli that were presented left vs. right of the fixation (V1) or activity for task vs. fixation (IPS). We used this statistic to rank the voxels within each ROI and selected voxels (500 for visual areas; 200 voxels for IPS) with the higher t-score to include in the MVPA, as classification accuracy saturated across all participants for these voxel pattern sizes in the corresponding regions (Figure S4). We used the same number of voxels (i.e. 200 voxels) when comparing data between V1 and IPS and for the informational connectivity analysis. This voxel selection procedure ensured that comparisons of MVPA accuracy could not be confounded by varying number of voxels across participants. We then extracted mean normalized fMRI responses between 4^th^ to 8^th^ TR (i.e. 6.18 – 14.42 s) after block onset for this pattern of voxels per ROI, participant and test session. We trained a linear classifier using LIBSVM (http://www.csie.ntu.edu.tw/~cjlin/libsvm/) implemented in MATLAB to discriminate: a) the trained from the control orientation, b) the untrained from the control orientation. As both the trained and untrained orientation differed equally from the control orientation (~55°), we hypothesized that differences in the accuracy between these two classification tasks would be due to training rather than stimulus differences. We computed classification accuracy using a leave-one-run-out cross-validation. That is, we divided the data set into training and test data with maximum 72 training patterns (for n = 7 participants with 8 runs) and 8 patterns for the test run. We averaged the classification accuracy across folds, separately for each test session. We used repeated-measures ANOVAs to assess differences in classification accuracy across conditions (orientation × session). Similar to the MPI for behavioral data, we defined the MPI for decoding accuracy as (post-test accuracy – pre-test accuracy) / pre-test accuracy × 100%).

Further, we performed a correlation-based pattern analysis^37^ to consolidate our MVPA results. In the correlation-based pattern analysis, the data and voxels used were identical to those used in the MVPA analysis. We divided the data set into training and test data and performed a leave-one-run-out cross-validation. For each dataset, we calculated the mean response of each orientation for each voxel. We then calculated the Spearman correlation across voxels and transformed the correlation coefficients using Fisher’s z-transform. We hypothesized that the correlation coefficient would be higher for data from the training and test set that related to the same orientation (i.e. trained-trained orientation) than different orientations (i.e. trained-control orientation). We used the difference between the same and different orientations to index the information contained in each ROI. We used repeated-measures ANOVAs to examine differences across conditions (orientation × session).

#### Informational Connectivity analysis

We used Informational Connectivity (IC) to identify layers that share synchronized discriminability of activity related to stimulus-specific multi-voxel pattern information^36,69,70^. We examined intercortical IC based on shared changes (fluctuations) in pattern discriminability over time, as this approach has been shown to be more sensitive than univariate functional connectivity. To track the flow of multivariate information across time (i.e. across blocks), we measured the fluctuations (covariance) in MVPA discriminability by calculating distance information from the classification hyperplane (Figure 7A). In particular, we selected 200 voxels with the higher t-score and used the same multivoxel training vs. test patterns as in the MVPA analysis. For each ROI and layer, we extracted distance information for the test data per block from the trained classifiers. We calculated layer-specific connectivity by partial Spearman correlation between the fold-wise distance of different layers; that is, for a given layer, we regressed out the distance information from other layers within each ROI. We transformed the correlation coefficients using Fisher’s z-transform and conducted repeated measures ANOVA to compare across conditions.

## Acknowledgements

We would like to thank Valentyna Chernova and Cher Zhou for help with the analysis. We are grateful to Cong Yu and Sheng Li for their helpful comments and suggestions on this work. This work was supported by grants to Z.K. from the Biotechnology and Biological Sciences Research Council (H012508 and BB/P021255/1), the Wellcome Trust (205067/Z/16/Z) and European Union’s Horizon 2020 research and innovation programme under the Marie Skłodowska Curie grant agreement No 840271. The 7T MRI was funded by the MRC (MR/M008983/1) and the NIHR Cambridge Biomedical Research Centre. C.T.R. is funded by the Wellcome Trust and the Royal Society (098436/Z/12/B).

## Author contributions

K.J. and Z.K. conceived the project and designed the experiments. V.K., C.R., C.T.R., G.W. and R.G. developed the GE EPI protocol used in the study. K.J. and E.Z. performed the experiments. K.J., E.Z, N.R.G., and A.N. developed the analysis pipeline and analyzed the data. K.J. and Z.K. wrote the manuscript.

## Supplementary Material

**Figure S1.**
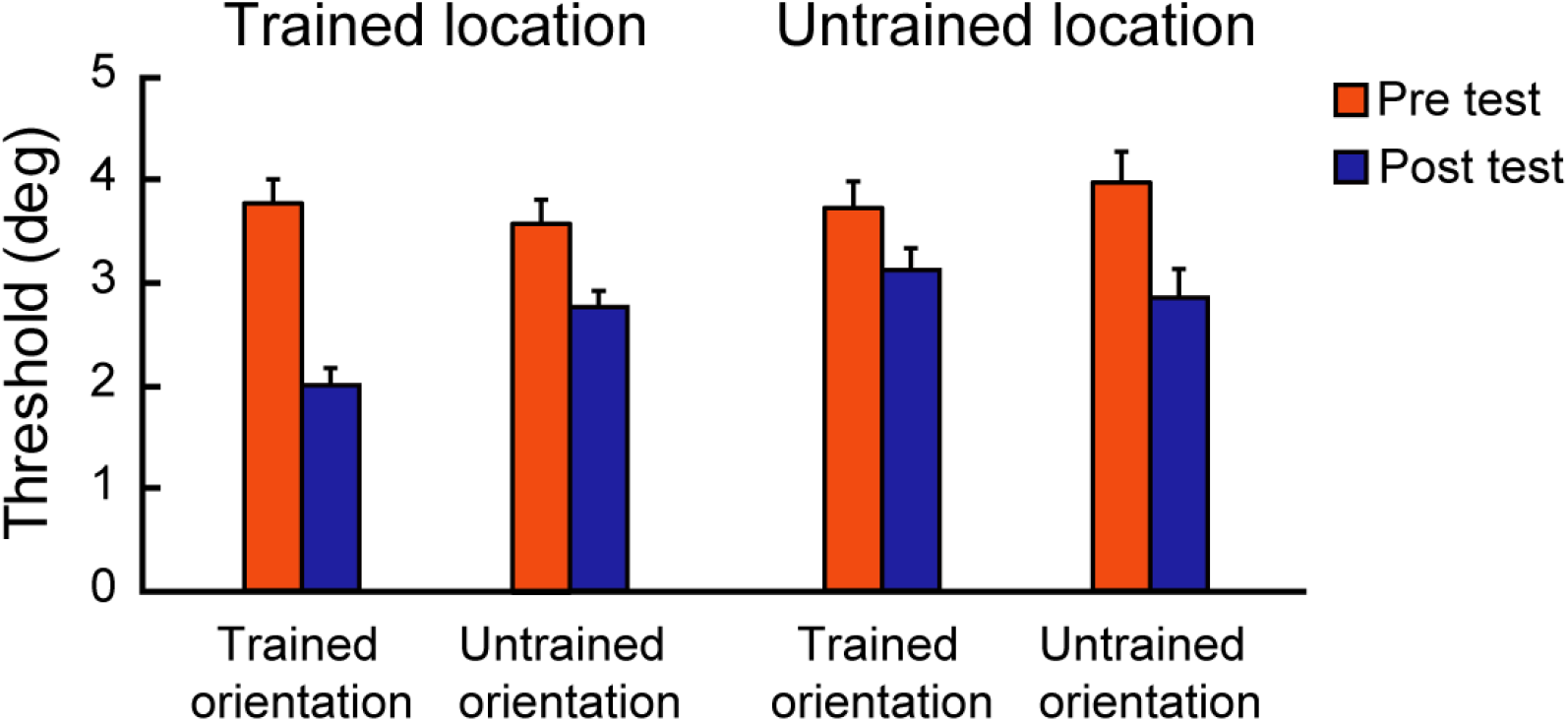
Behavioral results. Mean performance across participants before and after training (pre-, post-test) at ~79.4% threshold for the trained and untrained orientations presented at the trained and untrained locations. Error bars indicate standard error of the mean across participants. We observed learning specificity for the trained orientation at the trained location, as indicated by a significant orientation × location × session interaction (repeated measures ANOVA, F(1,12) = 11.858, *p* = 0.005) and a significant orientation × session interaction (F(1,12) = 21.551, *p* = 0.001) at the trained, but not the untrained (F(1,12) = 3.093, *p* = 0.104) location. Post-hoc comparisons at the trained location showed significantly lower threshold for the trained than the untrained orientation after (t(12) = −5.208, *p* < 0.001), but not before (t(12) = 1.264, *p* = 0.230) training. In contrast, no significant differences between the trained and the untrained orientations were observed at the untrained location (pre-test session: t(12) = −1.203, *p* = 0.252, post-test session: t(12) = 1.066, *p* = 0.308).

**Figure S2.**
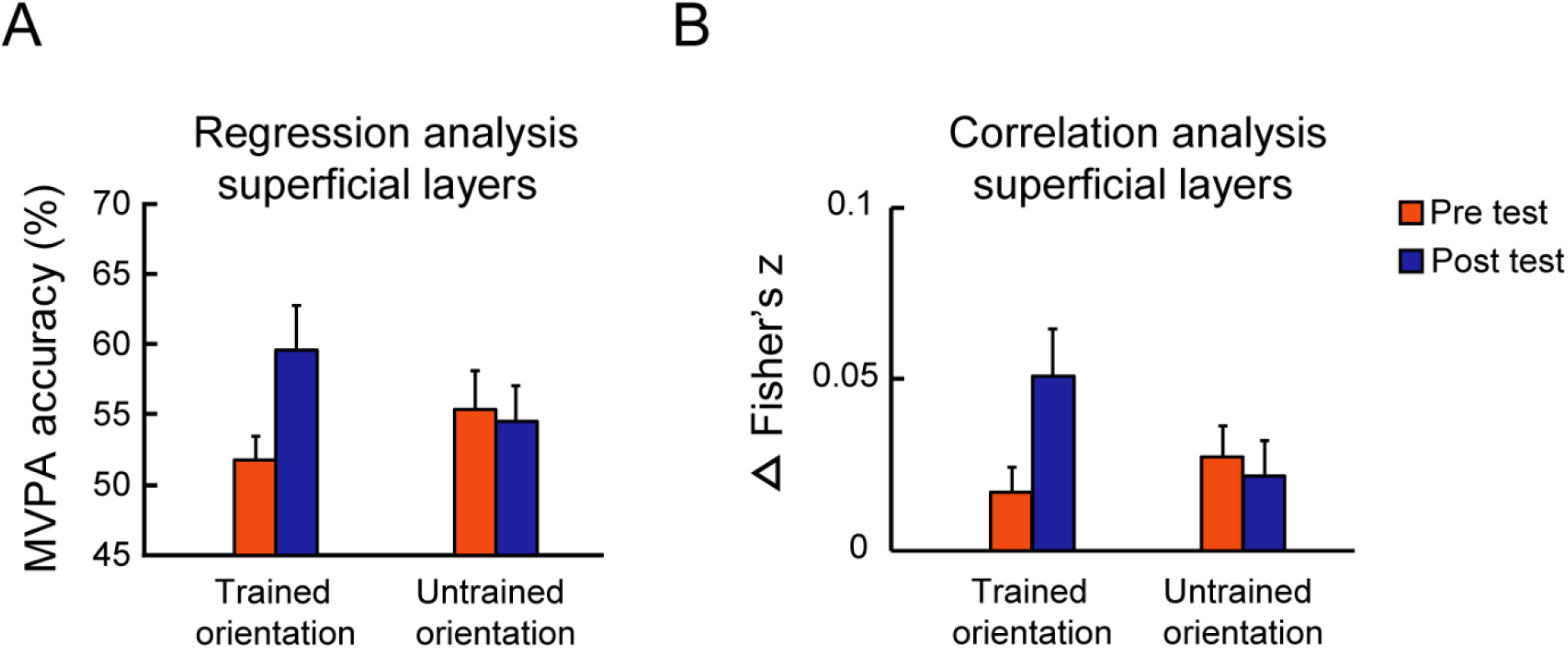
Control analyses. (A) MVPA accuracy before and after training for the trained and untrained orientations presented at the trained location in superficial V1 layers after regressing out the signal from the adjacent voxels in middle layers. (B) Correlation-based pattern analysis. Correlation differences (correlation of mean normalized BOLD across voxels for the same orientation minus normalized BOLD for different orientations) for the trained and untrained orientations presented at the trained location in superficial V1 layers. Error bars indicate standard error of the mean across participants.

**Figure S3.**
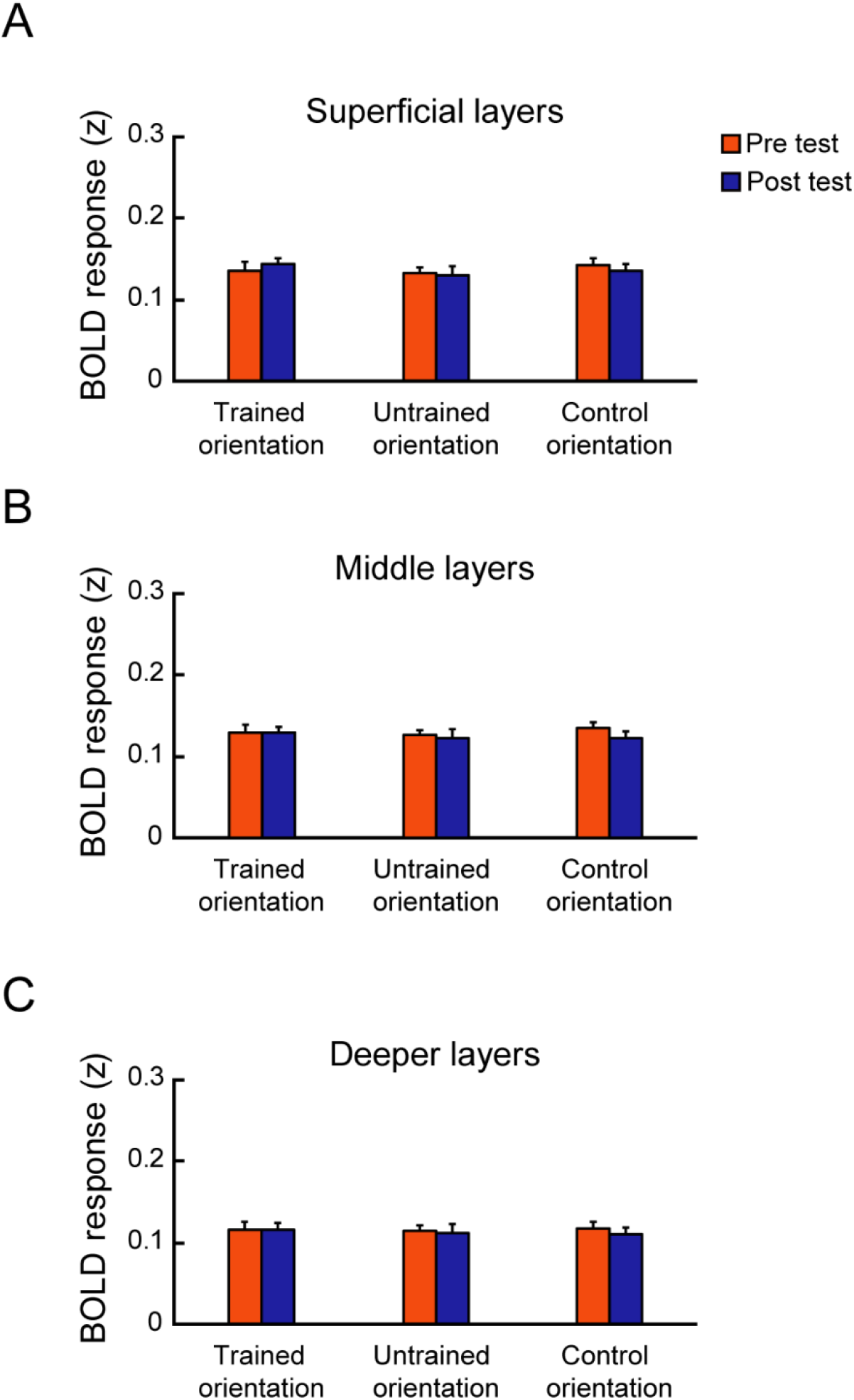
Univariate fMRI analysis. Mean normalized BOLD before and after training for the trained, untrained and control orientations presented at the trained location across V1 layers. Error bars indicate standard error of the mean across participants.

**Figure S4.**
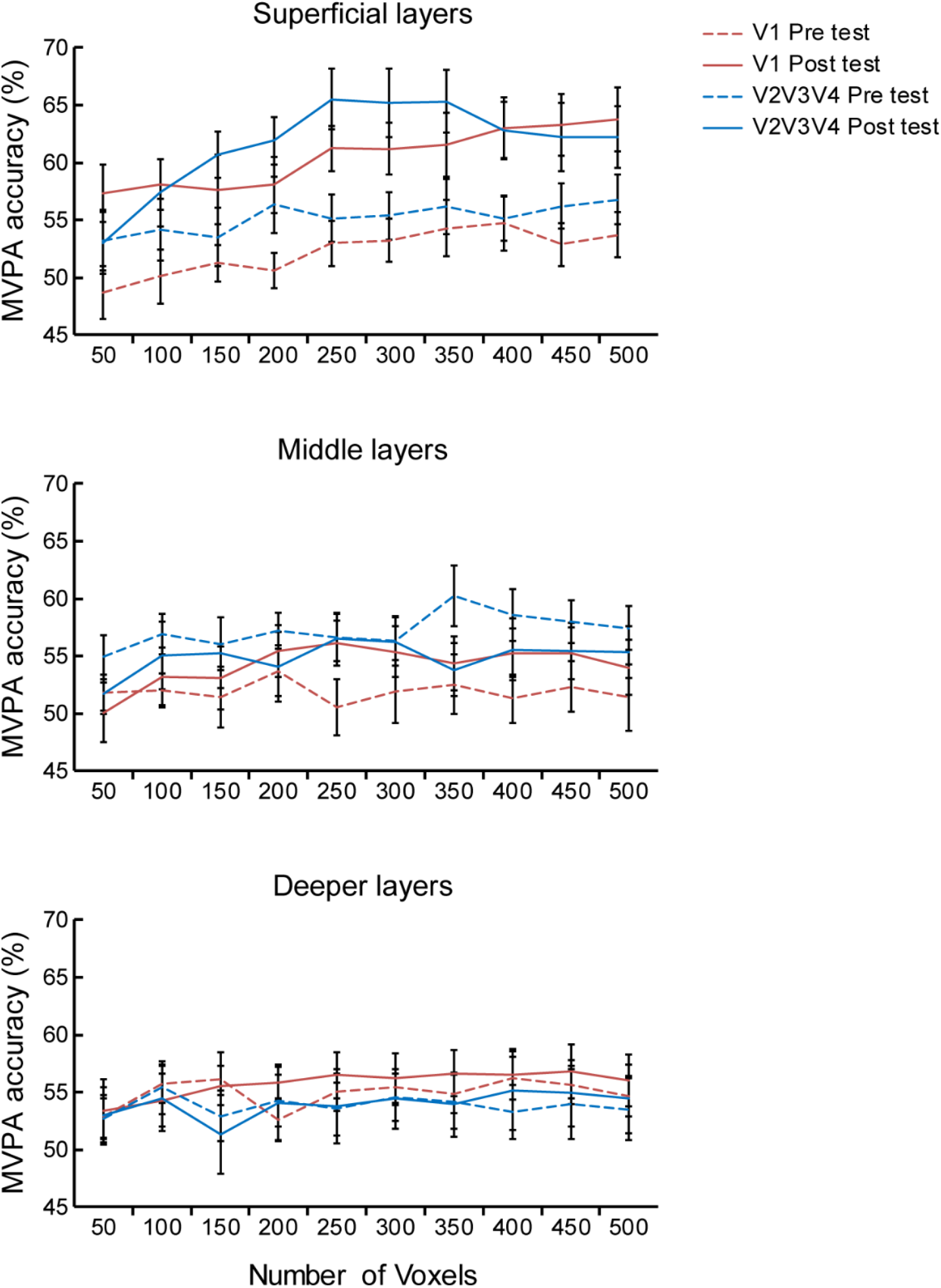
MVPA before and after training across visual areas. MVPA accuracy before and after training for different voxel patterns (from 100-500 voxels) for the trained orientation presented at the trained location across layers of V1, V2, V3, V4. We observed similar learning-dependent changes in superficial layers across visual areas. Two-way repeated measures ANOVAs (ROI × session) showed a significant main effect of session for pattern size of 200 (F(1,12) = 7.159, *p* = 0.020), 300 (F(1,12) = 7.751, *p* = 0.017), 400 (F(1,12) = 6.006, *p* = 0.031) voxels, and a trend for pattern size of 100 (F(1,12) = 3.743, *p* = 0.077) and 500 (F(1,12) = 4.587, *p* = 0.053) voxels. We did not observe any significant ROI × session interaction (all *ps* > 0.269). Further, we did not observe any significant differences in MVPA accuracy before vs. after training in middle nor deeper layers (all *ps* > 0.539). Error bars indicate standard error of the mean across participants.

## References

1. Gilbert, C. D., Sigman, M. & Crist, R. E. The Neural Basis of Perceptual Learning. Neuron 31, 681–697 (2001).

2. Gilbert, C. D. & Li, W. Adult Visual Cortical Plasticity. Neuron 75, 250–264 (2012).

3. Hooks, B. M. & Chen, C. Circuitry Underlying Experience-Dependent Plasticity in the Mouse Visual System. Neuron 106, 21–36 (2020).

4. Sagi, D. & Tanne, D. Perceptual learning: learning to see. Curr. Opin. Neurobiol. 4, 195–199 (1994).

5. Dosher, B. & Lu, Z.-L. Visual Perceptual Learning and Models. Annu. Rev. Vis. Sci. 3, 343–363 (2017).

6. Law, C. T. & Gold, J. I. Shared mechanisms of perceptual learning and decision making. Top. Cogn. Sci. 2, 226–238 (2010).

7. Ahissar, M. & Hochstein, S. Task difficulty and the specificity of perceptual learning. Nature 387, 401–406 (1997).

8. Schoups, A. A., Vogels, R. & Orban, G. a. Human perceptual learning in identifying the oblique orientation: retinotopy, orientation specificity and monocularity. J. Physiol. 483, 797–810 (1995).

9. Schoups, A., Vogels, R., Qian, N. & Orban, G. Practising orientation identification improves orientation coding in V1 neurons. Nature 412, 549–553 (2001).

10. Yan, Y. et al. Perceptual training continuously refines neuronal population codes in primary visual cortex. Nat. Neurosci. 17, 1380–1389 (2014).

11. Bao, M., Yang, L., Rios, C., He, B. & Engel, S. a. Perceptual Learning Increases the Strength of the Earliest Signals in Visual Cortex. J. Neurosci. 30, 15080–15084 (2010).

12. Jehee, J. F. M., Ling, S., Swisher, J. D., Bergen, R. S. Van & Tong, F. Perceptual learning selectively refines orientation representations in Early Visual Cortex. J. Neurosci. 32, 16747–16753 (2012).

13. Schwartz, S., Maquet, P. & Frith, C. Neural correlates of perceptual learning : A functional MRI study of visual texture discrimination. Proc. Natl. Acad. Sci. 99, 17137–17142 (2002).

14. Raiguel, S., Vogels, R., Mysore, S. G. & Orban, G. a. Learning to see the difference specifically alters the most informative V4 neurons. J. Neurosci. 26, 6589–6602 (2006).

15. Yang, T. & Maunsell, J. H. R. The Effect of Perceptual Learning on Neuronal Responses in Monkey Visual Area V4. J. Neurosci. 24, 1617–1626 (2004).

16. Kahnt, T., Grueschow, M., Speck, O. & Haynes, J. D. Perceptual Learning and Decision-Making in Human Medial Frontal Cortex. Neuron 70, 549–559 (2011).

17. Ahissar, M. & Hochstein, S. The reverse hierarchy theory of visual perceptual learning. Trends Cogn. Sci. 8, 457–464 (2004).

18. Goense, J., Bohraus, Y. & Logothetis, N. K. fMRI at high spatial resolution: implications for BOLD-models. Front. Comput. Neurosci. 10, (2016).

19. RockLand, K. S. & Pandya, D. N. Laminar origins and terminations of cortical connections of the occipital lobe in the rhesus monkey. Brain Res. 179, 3–20 (1979).

20. Markov, N. T. et al. Anatomy of Hierarchy: Feedforward and Feedback Pathways in Macaque Visual Cortex. J. Comp. Neurol. 522, 225–259 (2014).

21. Larkum, M. E., Petro, L. S., Sachdev, R. N. S. & Muckli, L. A Perspective on Cortical Layering and Layer-Spanning Neuronal Elements. Front. Neuroanat. 12, (2018).

22. Self, M. W., van Kerkoerle, T., Goebel, R. & Roelfsema, P. R. Benchmarking laminar fMRI: Neuronal spiking and synaptic activity during top-down and bottom-up processing in the different layers of cortex. Neuroimage 197, 806–817 (2019).

23. Gilbert, C. D. & Wiesel, T. Clustered intrinsic connections in cat visual cortex. J. Neurosci. 3, 1116–1133 (1983).

24. Douglas, R. J. & Martin, K. A. C. Recurrent neuronal circuits in the neocortex. Curr. Biol. 17, 496–500 (2007).

25. Schwabe, L. & Obermayer, K. Adaptivity of tuning functions in a generic recurrent network model of a cortical hypercolumn. J. Neurosci. 25, 3323–3332 (2005).

26. Self, M. W., van Kerkoerle, T., Supèr, H. & Roelfsema, P. R. Distinct roles of the cortical layers of area V1 in figure-ground segregation. Curr. Biol. 23, 2121–2129 (2013).

27. Buffalo, E. A., Fries, P., Landman, R., Buschman, T. J. & Desimone, R. Laminar differences in gamma and alpha coherence in the ventral stream. Proc. Natl. Acad. Sci. 108, 11262–11267 (2011).

28. Martino, F. De et al. Frequency preference and attention effects across cortical depths in the human primary auditory cortex. Proc. Natl. Acad. Sci. 112, 16036–16041 (2015).

29. Lawrence, S. J. D., Norris, D. G. & De Lange, F. P. Dissociable laminar profiles of concurrent bottom-up and top-down modulation in the human visual cortex. Elife 8, 1–17 (2019).

30. Hung, S. & Seitz, A. R. Prolonged Training at Threshold Promotes Robust Retinotopic Specificity in Perceptual Learning. J. Neurosci. 34, 8423–8431 (2014).

31. Zhang, J. Y. et al. Rule-based learning explains visual perceptual learning and its specificity and transfer. J. Neurosci. 30, 12323–12328 (2010).

32. Uǧurbil, K., Toth, L. & Kim, D. S. How accurate is magnetic resonance imaging of brain function? Trends Neurosci. 26, 108–114 (2003).

33. Kay, K. et al. A critical assessment of data quality and venous effects in sub-millimeter fMRI. Neuroimage 189, 847–869 (2019).

34. Vizioli, L. et al. Multivoxel Pattern of Blood Oxygen Level Dependent Activity can be sensitive to stimulus specific fine scale responses. bioRxiv (2019).

35. Kok, P. et al. Selective Activation of the Deep Layers of the Human Primary Visual Cortex by Top-Down Feedback. Curr. Biol. 26, 371–376 (2016).

36. Koster, R., Chadwick, M. J., Chen, Y. & Kumaran, D. Big-Loop Recurrence within the Hippocampal System Supports Integration of Information across Episodes. Neuron 99, 1342–1354 (2018).

37. Haxby, J. V. et al. Distributed and overlapping representations of faces and objects in ventral temporal cortex. Science (80-.). 293, 2425–2430 (2001).

38. Chen, N. et al. Sharpened cortical tuning and enhanced cortico-cortical communication contribute to the long-term neural mechanisms of visual motion perceptual learning. Neuroimage 115, 17–29 (2015).

39. Byers, A. & Serences, J. T. Exploring the relationship between perceptual learning and top-down attentional control. Vision Res. 74, 30–39 (2012).

40. Gilbert, C. D., Li, W. & Piech, V. Perceptual learning and adult cortical plasticity. J. Physiol. 587, 2743–51 (2009).

41. Zhang, J., Meeson, A., Welchman, A. E. & Kourtzi, Z. Learning alters the tuning of functional magnetic resonance imaging patterns for visual forms. J. Neurosci. 30, 14127–14133 (2010).

42. Furmanski, C. S., Schluppeck, D. & Engel, S. A. Learning Strengthens the Response of Primary Visual Cortex to Simple Patterns. Curr. Biol. 14, 573–578 (2004).

43. Mukai, I. et al. Activations in visual and attention-related areas predict and correlate with the degree of perceptual learning. J. Neurosci. 27, 11401–11411 (2007).

44. Ringach, D. L., Shapley, R. M. & Hawken, M. J. Orientation selectivity in macaque V1: diversity and laminar dependence. J. Neurosci. 22, 5639–5651 (2002).

45. Gau, R., Bazin, P.-L., Trampel, R., Turner, R. & Noppeney, U. Resolving multisensory and attentional influences across cortical depth in sensory cortices. Elife 9, (2020).

46. Muckli, L. et al. Contextual Feedback to Superficial Layers of V1. Curr. Biol. 25, 2690–2695 (2015).

47. Chang, D. H. F., Mevorach, C., Kourtzi, Z. & Welchman, A. E. Training transfers the limits on perception from parietal to ventral cortex. Curr. Biol. 24, 2445–2450 (2014).

48. Malach, R., Amir, Y., Harel, M. & Grinvald, A. Relationship between intrinsic connections and functional architecture revealed by optical imaging and in vivo targeted biocytin injections in primate striate cortex. Proc. Natl Acad. Sci. USA 90, 10469–10473 (1993).

49. Teich, A. F. & Qian, N. Learning and adaptation in a recurrent model of V1 orientation selectivity. J. Neurophysiol. 89, 2086–2100 (2003).

50. Huber, L., Uludağ, K. & Möller, H. E. Non-BOLD contrast for laminar fMRI in humans: CBF, CBV, and CMRO2. Neuroimage 197, 742–760 (2019).

51. Watanabe, T. & Sasaki, Y. Perceptual Learning: Toward a Comprehensive Theory. Annu. Rev. Psychol. 66, 1–25 (2015).

52. Lewis, C. M., Baldassarre, A., Committeri, G., Romani, G. L. & Corbetta, M. Learning sculpts the spontaneous activity of the resting human brain. Proc. Natl. Acad. Sci. 106, 17558–17563 (2009).

53. Van Essen, D. C. & Felleman, D. J. Distributed Hierarchical Processing in the Primate Cerebral Cortex. Cereb. Cortex 1, 1–47 (1991).

54. Pelli, D. G. The VideoToolbox software for visual psychophysics: transforming numbers into movies. Spat. Vis. 10, 437–442 (1997).

55. Brainard, D. H. The Psychophysics Toolbox. Spat. Vis. 10, 433–436 (1997).

56. Moeller, S. et al. Multiband multislice GE-EPI at 7 tesla, with 16-fold acceleration using partial parallel imaging with application to high spatial and temporal whole-brain FMRI. Magn. Reson. Med. 63, 1144–1153 (2010).

57. Waehnert, M. D. et al. Anatomically motivated modeling of cortical laminae. Neuroimage 93, 210–220 (2014).

58. Kemper, V. G., De Martino, F., Emmerling, T. C., Yacoub, E. & Goebel, R. High resolution data analysis strategies for mesoscale human functional MRI at 7 and 9.4 T. Neuroimage 164, 48–58 (2018).

59. Greve, D. N. & Fischl, B. Accurate and robust brain image alignment using boundary-based registration. Neuroimage 48, 63–72 (2009).

60. Engel, S. a, Wandell, B. a & Glover, G. G. Retintopic organization in human visual cortex and the spatial precision of functional MRI. Cereb. Cortex 7, 181–192 (1997).

61. Sereno, M. I. et al. Borders of Multiple Visual Areas In Humans Revealed By Functional MRI. Science (80-.). 268, 889–893 (1995).

62. Wang, L., Mruczek, R. E. B., Arcaro, M. J. & Kastner, S. Probabilistic maps of visual topography in human cortex. Cereb. Cortex 25, 3911–3931 (2015).

63. Uludaǧ, K., Müller-Bierl, B. & Uǧurbil, K. An integrative model for neuronal activity-induced signal changes for gradient and spin echo functional imaging. Neuroimage 48, 150–165 (2009).

64. Yacoub, E., Van De Moortele, P. F., Shmuel, A. & Uǧurbil, K. Signal and noise characteristics of Hahn SE and GE BOLD fMRI at 7 T in humans. Neuroimage 24, 738–750 (2005).

65. Duvernoy, H. M., Delon, S. & Vannson, J. L. Cortical blood vessels of the human brain. Brain Res. Bull. 7, 519–579 (1981).

66. Olman, C. A., Inati, S. & Heeger, D. J. The effect of large veins on spatial localization with GE BOLD at 3 T: Displacement, not blurring. Neuroimage 34, 1126–1135 (2007).

67. Polimeni, J. R., Fischl, B., Greve, D. N. & Wald, L. L. Laminar analysis of 7 T BOLD using an imposed spatial activation pattern in human V1. Neuroimage 52, 1334–1346 (2010).

68. Kashyap, S., Ivanov, D., Havlicek, M., Poser, B. A. & Uludağ, K. Impact of acquisition and analysis strategies on cortical depth-dependent fMRI. Neuroimage 168, 332–344 (2018).

69. Coutanche, M. N. & Thompson-Schill, S. L. Using informational connectivity to measure the synchronous emergence of fMRI multi-voxel information across time. J. Vis. Exp. 1–7 (2014) doi:10.3791/51226.

70. Anzellotti, S. & Coutanche, M. N. Beyond Functional Connectivity: Investigating Networks of Multivariate Representations. Trends Cogn. Sci. 22, 258–269 (2018).

